# GlycansToGraphs: visualizing and simplifying complex mass spectra

**DOI:** 10.1101/2024.01.31.578148

**Authors:** R. Bonner, C. Jacquet, G. Hopfgartner

## Abstract

We describe a software tool, GlycansToGraphs, used to identify and visualize relationships between masses in MS or MS/MS spectra and illustrate its application to data dependent acquisition (DDA) spectra of glycopeptides. The software is written in python 3.11 and uses the ‘streamlit’ package (1.29) to generate a User Interface (UI) in a web browser. It uses simple text input files, does not require databases (glycan or protein) and the user can define any mass difference that the software searches for in an unbiased manner. Located mass differences generate graphs with edges that correspond to the relationships between nodes (masses) and collections of connected nodes, known as components, which can be analyzed to extract sequences of relationships.

## 2. Introduction

Interpretation of the MS and tandem MS (MS/MS) spectra from MS and LCMS using electrospray for the analysis of metabolites or peptides is complicated by a variety of factors. Firstly, many analytes are inherently heterogeneous, for example, post-translational protein glycosylation can introduce heterogeneity at the macro level (different glycans, different glycosites) and micro level (glycan isomers) producing chemically similar species(1). Secondly, the analysis method can introduce artifacts. Proteins are often analyzed by LC-MS/MS of tryptic digests treated to remove disulphide links which can produce missed cleavages and off-target (i.e. non Cys) derivatives. Thirdly, mass spectral annotation is complicated because individual analytes can produce large numbers of different ions(2, 3). For example, Farke *et al*. (4) recently described the flow injection analysis (FIA) of 160 compounds individually spiked into an E. coli extract and reported that on average 68 features were detected for each analyte with 11 compounds producing more than 100 features and one, glycerol 3-phsophate, producing an astonishing 5464. Finally, complete characterization requires combining the results of different analytical techniques such as: ion polarity, LC column and digestion enzymes, ion mobility, and various ion activation methods (CAD, UVPD, ExD, EThcD).

Depending on the system the observed ions may be expected (isotopes, different charge, different charge carriers), variable (known but unpredictable, e.g. missed cleavages, off-target derivatization intended for Cys-Cys or unusual adducts such as Al, Ba, Fe) or unexpected (reactions with background ions or co-eluting species in LC/MS). Several well-known data processing steps are available to initially simplify interpretation, especially de-isotoping to remove 13C peaks and deconvolution to combine related ions with different charge states, but these frequently leave unidentified species. However, molecules of interest are often related by a few species with known masses (e.g. the individual sugars in glycosylation, or amino acids in peptides). Thus a valuable step is the unbiased identification of groups of ions connected by these expected mass differences and the generation and visualization of the corresponding networks. The termini of these networks can indicate the root masses of interest or point to unknown processes that must be identified and incorporated in further processing cycles. However, while the complex patterns can explain the observed spectrum, determining the roots of the patterns is essential to identifying the analytes of interest.

Here we describe a software tool that uses text input files from MS or MS/MS spectra and lists of user-specified mass differences to identify and visualize relationships between masses and illustrate its application to data dependent acquisition (DDA) spectra of glycopeptides. Essentially the tool filters complex spectra by generating networks of ions related by specific masses, in this case glycans, but generic values corresponding to adducts, fragments or any mass of interest can also be used.

## 3. Software description

GlycansToGraphs (G2G) uses directed graphs defined by a set of ‘nodes’ connected by ‘edges’. A node is a mass, with the underlying peak as an attribute, and an edge exists between two nodes if the mass difference corresponds to a target mass within a user-specified tolerance. The edge direction indicates the parent and child nodes with the lower mass being the parent unless the target mass corresponds to a neutral loss specified as a negative value. If desired all edges can be treated as losses, i.e. the higher mass is always the parent. To construct the graph the program considers each mass once, searches for relative target masses and, if one is found, records both masses in a unique set of nodes and creates the edge between them. Paths longer than one target mass are found automatically since if A -> B -> C, A -> B is added when A is considered and B -> C when B is considered. Unconnected masses are currently ignored.

Once the graph is constructed algorithms provided by the python packages networkX and graphviz are used for layout and analysis. The graph can be divided into ‘components’ that are independent collections of connected nodes, and sources (nodes with no parents) and sinks (nodes with no children) found. These end points are the root masses to be explained and paths connecting them can be identified to determine sequences. Sequences are stored as a list of individual members in the order detected, e.g. N.N.H and N.H.N, and in a canonical form that records the composition, i.e. H.N2 for these. This allows rapid comparison while retaining the order of elements for further analysis.

Targets have the several attributes in addition to the mass: a name (e.g. ‘Hex’), a symbol used to label the edges and in sequences (e.g. ‘H’), a colour for the edge in the displayed graph, and a boolean value indicating whether this mass is enabled by default in the UI. Values specified in a configuration file are automatically included in the user interface, but the user can also specify one or more element symbols or enter arbitrary masses as text strings separated by semi-colons and optionally providing a label and a colour. The latter is useful for interactively observing the effects of arbitrary masses on the network. Allowed elements are from a limited set (currently O, Na, K, Mg, Al, Ca, Fe3, Fe2, Ni, Co, Zr, Ba) that generate ‘effective adduct masses’ {Bonner, 2022 #2} i.e. considering the oxidation state of the element, thus Fe can be II (Fe2) or III (Fe3).

Peak lists are generally independent but it can be useful to process multiple spectra, for example, glycopeptide MSMS spectra from peaks with the same mass but different charges. Similarly, extensive heterogeneity can generate closely eluting spectra that are very similar. Thus the software accepts one or more mass spectra supplied as text files containing m/z, intensity, charge state (z) and monoisotopic status. Spectra are converted to mass using z if provided otherwise 1 is assumed. Peaks with charge of zero, indicating that no isotope information was detected, can be dropped or converted assuming they are singly charged. Peaks that are close in mass can be combined which simplifies visualization for spectra that contain multiply charged ions of the same molecule, although it can be useful to observe that a mass is derived from more than one m/z value. Note that converting the spectra to mass allows different charge states and polarities to be directly compared.

The graph can be based on all peaks or a subset selected based on intensity and monoisotopic status, if available. Using only monoisotopic peaks simplifies the graph by removing the multiple edges generated by isotopes but relies on the peak finding software accurately determining these peaks. Peaks from different files can be treated independently or combined (‘consolidated’) which can be useful, for example, when precursor ions with different charges generate different fragments. After generating the graph the program determines the components and summarizes them as a text histogram based on the size (number of nodes) and their intensity sum. Since graphs often contain many components with one or two edges, the user can provide a threshold to filter those displayed. If the number of nodes exceeds a threshold value the graph is simplified by hiding the edge labels and reducing the displayed precision of the masses.

The components are also summarized in a table that includes size, intensity sum and the highest and lowest masses and the user can chose to analyze one or all of these further by determining and tabulating the sources, sinks and connecting paths. The results can be saved as a CSV file for further analysis elsewhere.

## 4. Data Processing

Processing can be applied to one or more data files and proceeds in several steps:

### 4.1 Conversion to mass

- Peak lists are exported from PeakView software (Sciex) via ‘Export peaks as text..’ which produces an extensive table that includes m/z, charge, mass, monoisotopic, height, etc.
- Some peaks may have a charge of zero if no obvious isotope peaks (i.e. with expected spacing) were observed. These can be ignored or treated as singly charged (the most likely scenario for isolated ions)
- Since the charge is identified, measured *m/z* values can be converted to mass by assuming the charge is due to protons, i.e. *m/z* = (M + nH+)/n => M = n(m/z - H+)
- There are options to merge and consolidate the peak lists. Since we are dealing with different charge states, and potentially different files, the same mass may be present several times. ‘Merge peaks’ combines peaks that are close in mass within a file while ‘Consolidate’ will combine nodes between files. For merged/consolidated peaks, ‘z’ becomes the number of merged nodes rather than charge.

### 4.2 Graph

- A graph consists of nodes (peaks) connected by edges (relationships)
- Nodes corresponds to masses and are stored with attributes, currently the corresponding peak and file identification number
- Edges correspond to known mass differences, taken from a list of target ‘molecules’, specified as name, symbol, expected mass difference and display colour.
- The main graph is directed, i.e. each edge has a direction, here from the source node to the target node. With the exception of neutral losses such as water, the source node is the lower mass, i.e. if H represents hexose and is the mass difference between A and B, then A -> B means A + H = B.
- There is an option to treat all differences as losses, i.e. the higher mass is always the source.
- The edge label includes the symbol of the associated molecule and, optionally, the mass error in mmu (i.e. measured mass difference minus the expected value).

### 4.3 Graph generation

- Graphs can be generated from all peaks or only the monoisotopic.

Graphs are generated by comparing the mass difference between all pairs of peaks to a list of target masses (molecules) using a tolerance that is specified as the greater of a value in mmu or ppm. Note: since we are considering masses the values, and hence ppm tolerance, can be large.

- When a match is found the two peaks are stored as nodes, if not already stored, and an edge is created between them. Paths involving multiple masses are found automatically when a nodes is both a source and a target. Given the list of nodes and edges, the graph software will find the sequences.

### 4.4 Filtering

- Prior to graph generation the underlying list of peaks can be filtered in various ways including by charge state, intensity and monoisotopic status.

Charge state filtering is only available if peaks are not merged or consolidated

- Since the number of peaks/nodes can be large ‘Consolidate’ and ‘Mono only’ are the default settings.
- The final display can be filtered by removing ‘components’ (see below) for simplification.

### 4.5 Graph drawing

- Currently the graphs are drawn using the python packages networkX and graphviz, which maintain the graphs and calculate the node positions, and matplotlib for the final visualization.
- There are many algorithms for calculating the layout; we have found that graphviz/dot and graphviz/neato are the most useful here but others are available and can be explored.
- The graphing libraries contain many algorithms for searching graphs, finding paths and distances, etc. GlycansToGraphs uses ‘components’ and paths: components are sets of related nodes and paths are lists of nodes that connect two nodes. MS data often has many small components, i.e. two peaks related by a target mass but otherwise unconnected: these can be useful for explaining the spectrum but not for understanding the sequences and can be hidden.
- Comments
- In the final graphs, node colour is used to represent charge state or the number of underlying merged peaks, node size is related to sqrt(intensity), node shape indicates the data file (currently no legend and merged nodes will use have the symbol of the file with the lowest index number). Shapes used, in order, are ‘ovsd’ (= circle, downward pointing triangle, square, diamond).
- Edge colour is arbitrary but using different colors for the main targets (sugars, CAM) helps visualize important sequences, e.g. N.N.H.H for N-linked glycans. Edge labels show the symbol of the corresponding target and can also include the mass error in mmu.
- ‘On the fly’ molecules are black unless specified otherwise.
- Filtering, including removing some target molecules, is useful for simplifying the graphs and helps understand peak relationships/sequences. Some relationships are useful as additional node evidence and for determining the explained peaks, but do not contribute to the sequence knowledge. Adducts, ^13^C, small neutral losses (e.g. H_2_O) and sample prep artifacts (multiple CAM addition) are in this category.

### 4.6 Component analysis

- Visualizing components is valuable for showing the extent and complexity of the relationships, but interpretation generally requires reducing these to ‘sequences’, i.e. a list of the symbols (obtained from the edges). The user can also look for specific sequences, regardless of mass (e.g. N-glycans)

### Analysis proceeds as follows

- For each component:
- Find all sources (no inputs), sinks (no outputs) and paths between sources and sinks as sequences By default this includes all intermediate paths, i.e. the path H.H.A.N generates H, H.H, H.H.A and H.H.A.N, and will contain many duplicates. An option allows only the complete path to be stored.
- Remove exact duplicates, i.e. considering all columns (masses, members in order, canonical form) This leaves all forms that differ by at least one field, i.e. different masses or member order, but can be filtered further.
- Filter as required:
- ‘Show first canonical form’ - shows sequences that have the same ID, start and end mass, and canonical form, i.e. hides sequences that differ only in the order of the members.
- ‘Ignore masses’ - modifies ‘Show first canonical form’ to show single canonical form, regardless of mass, members, etc. Essentially this shows the unique sequence compositions present anywhere in the selected component(s).
- ‘Filter’ and ‘Filter position’ - search the ‘Members’ column for a specific pattern at the start, end or anywhere.
- The displayed sequence table can be saved to a csv file if required

## 5. User interface description

The software is written in python 3.11 and uses the ‘streamlit’ package (1.29) to generate a User Interface (UI) in a web browser as shown in Figure 2 (detailed description is provided in Supplemental Info Figure S1). Target masses specified in a Tom’s Obvious Minimal Language (TOML) text configuration file are incorporated in the user interface but, importantly, arbitrary masses can be entered to interactively observe the effect on the network. If the charge state is specified, *m/z* values are converted to mass assuming protonation or deprotonation. The tool can process all or specific charge states which can be useful since glycopeptide spectra often contain singly charged B fragment ions corresponding to the glycan and multiply charged Y ions that also contain the peptide backbone(5). The resulting masses that are close in mass can be combined if desired.

The UI is in two parts: 1) a sidebar that can be hidden (boxes A, B, C) where the input file, target molecules and low level processing parameters are specified, and 2) an area for the graph and related parameters (boxes D, E, F, G, H).

## 6. Example

A bovine Fetuin (Sigma) digest was prepared. Fetuin was solubilized at 1ug/uL, buffer exchanged with 6M urea and 50mM Tris-HCl and heated at 95°C for 15min. This was followed by successive incubations: 20 minutes at 60 °C with 5 mM dithiothreitol (Fluka), 15 minutes in the dark at 25 °C with 15 mM iodoacetamide (Sigma), and 12 hours at 37 °C with 1:50 trypsin (Promega) after which digestion was stopped with 1 % formic acid (FA). The digestion was lyophilized and further dissolved in H_2_O with 0.1%FA for a final concentration of 1 μg/μL of the initial protein.

5μL of sample were injected using a column switching set up for sample cleaning and preconcentration. Trapping was performed at 150 μL/min at 95/5 H_2_O with 0.01% trifluoroacetic acid (TFA)/ACN with 0.01%TFA on an OPTI-Guard C8 (1 mm × 14 mm, Optimize Technologies) at room temperature. Gradient separation was performed at 20uL/min on a C18 Halo column (100 mm × 0.5 mm I.D, particle size of 2.7 μM, Halo) at 45 °C. The gradient was 7%B from 0 to 3 minutes, from 7% to 80%B from 3 to 18 minutes, 80%B from 18 to 23 minutes, 7%B from 23 to 25 minutes where A was H_2_O with 0.1%FA and B was ACN with 0.1% FA.

Mass spectrometry experiments were performed on a ZenoTOF 7600 (Sciex) equipped with the OptiFlow ion source. Data dependent acquisition (DDA) was used switching after 25 spectra with the following criteria: ions greater than 200 Da, charge state from 2 to 10, and intensity greater than 100 counts. Previous precursor ions were excluded for 20 seconds after 10 occurrences. Parameters used were source temperature 100 °C, IonSpray Voltage Floating 4500 V, collision energy 40 V, collision energy spray 10 V.

The survey scan Total Ion Chromatogram from the LC-MS/MS analysis and a representative spectrum are presented in Figure 3 and the exported text data file (GlycoPep_RT7_9.txt) used in GlycansToGraphs (G2G) is shown in Table 1. The minimum format requirement for the text file is Mass/Charge, Height, Charge, Monoisotopic status. The spectrum contains singly charged ions corresponding to glycan fragments and combinations (366.1397 = HexNAc.Hex, 657.2356 = HexNAc.Hex.NeuNAc) and a series of doubly charged ions at higher m/z. The graphical output from G2G for this spectrum is presented in Figure 4 and shows the relationships (glycans A,N,H) between the matched masses. Two components are apparent. The source node with mass 1739.82 is derived from the doubly charged series and clearly corresponds to the beginning of the N-glycan core structure, i.e. NexNAc.HexNAc.Hex.Hex (N.N.H.H). Hence it is likely a peptide, in this case the fetuin tryptic peptide LCPDCPLLAPLNDSR with both cysteine residues carbamidomethylated. The second component, generated by the singly charged low mass ions, has two source nodes, 203.080 and 291.096, which correspond to HexNAc and NeuAc respectively, and hence reflects the structure of the glycan.

**Table 1:** Format of the exported MS/MS spectrum (file: GlycoPep_RT7_9.txt).

| Mass/Charge | Area | Height | Width | Width at 50% | Resolution | Charge | Monoisotopic | Mass (charge) | Mass/charge (charge) |
| --- | --- | --- | --- | --- | --- | --- | --- | --- | --- |
| 109.0286 | 0.63 | 103 | 0.01 | 0.0065 | 16820 | 0 | Yes | *108.0208 | *109.0286 |
| 121.0288 | 7.72 | 1190 | 0.03 | 0.0059 | 20522 | 1 | Yes | *120.0209 (1) | *121.0288 (1) |
| 126.0552 | 10.64 | 1712 | 0.02 | 0.0058 | 21619 | 1 | Yes | *125.0474 (1) | *126.0552 (1) |
| 138.0550 | 63.31 | 10475 | 0.03 | 0.0057 | 24156 | 1 | Yes | *137.0471 (1) | *138.0550 (1) |
| 139.0581 | 5.08 | 610 | 0.05 | 0.0065 | 21267 | 1 | No | 138.0503 (1) | 139.0581 (1) |
| ..... |  |  |  |  |  |  |  |  |  |
| 635.3271 | 7.28 | 395 | 0.05 | 0.0177 | 35974 | 2 | Yes | *1268.6385 (2) | *635.3271 (2) |
| 635.8318 | 4.67 | 325 | 0.06 | 0.0104 | 60975 | 2 | No | 1269.6479 (2) | 635.8318 (2) |
| ..... |  |  |  |  |  |  |  |  |  |
| 1830.2679 | 10.83 | 185 | 0.16 | 0.0397 | 46151 | 2 | No | 3658.5202 (2) | 1830.2679 (2) |
| 1830.7447 | 5.79 | 134 | 0.12 | 0.0689 | 26572 | 2 | No | 3659.4737 (2) | 1830.7447 (2) |
| 1943.9146 | 25.63 | 370 | 0.16 | 0.0608 | 31983 | 1 | Yes | *1942.9068 (1) | *1943.9146 (1) |
| 1944.9220 | 27.84 | 486 | 0.16 | 0.0489 | 39802 | 1 | No | 1943.9142 (1) | 1944.9220 (1) |
| 1945.9157 | 15.53 | 204 | 0.23 | 0.0693 | 28080 | 1 | No | 1944.9078 (1) | 1945.9157 (1) |

**Figure 1.**
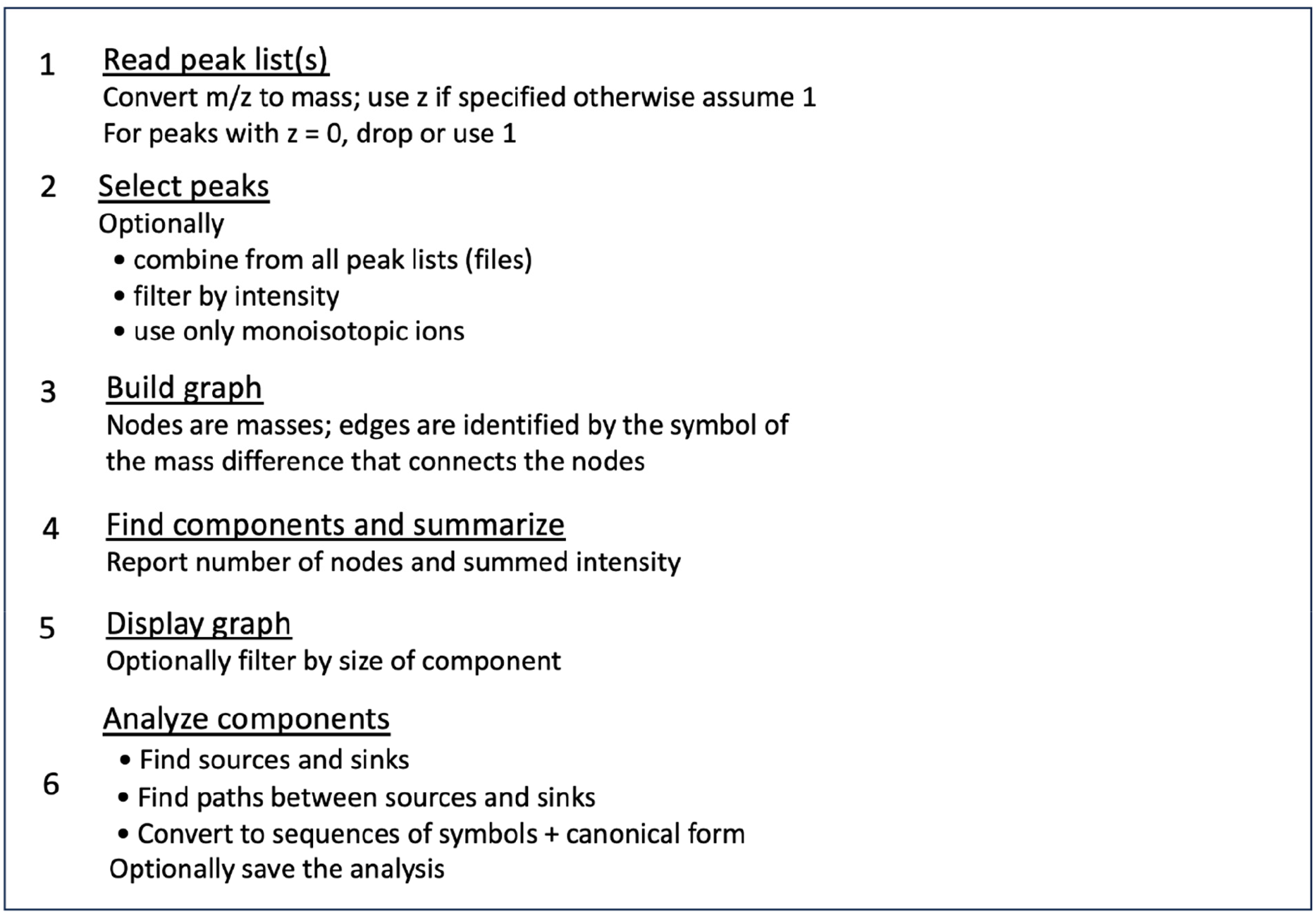
Processing workflow summary

**Figure 2.**
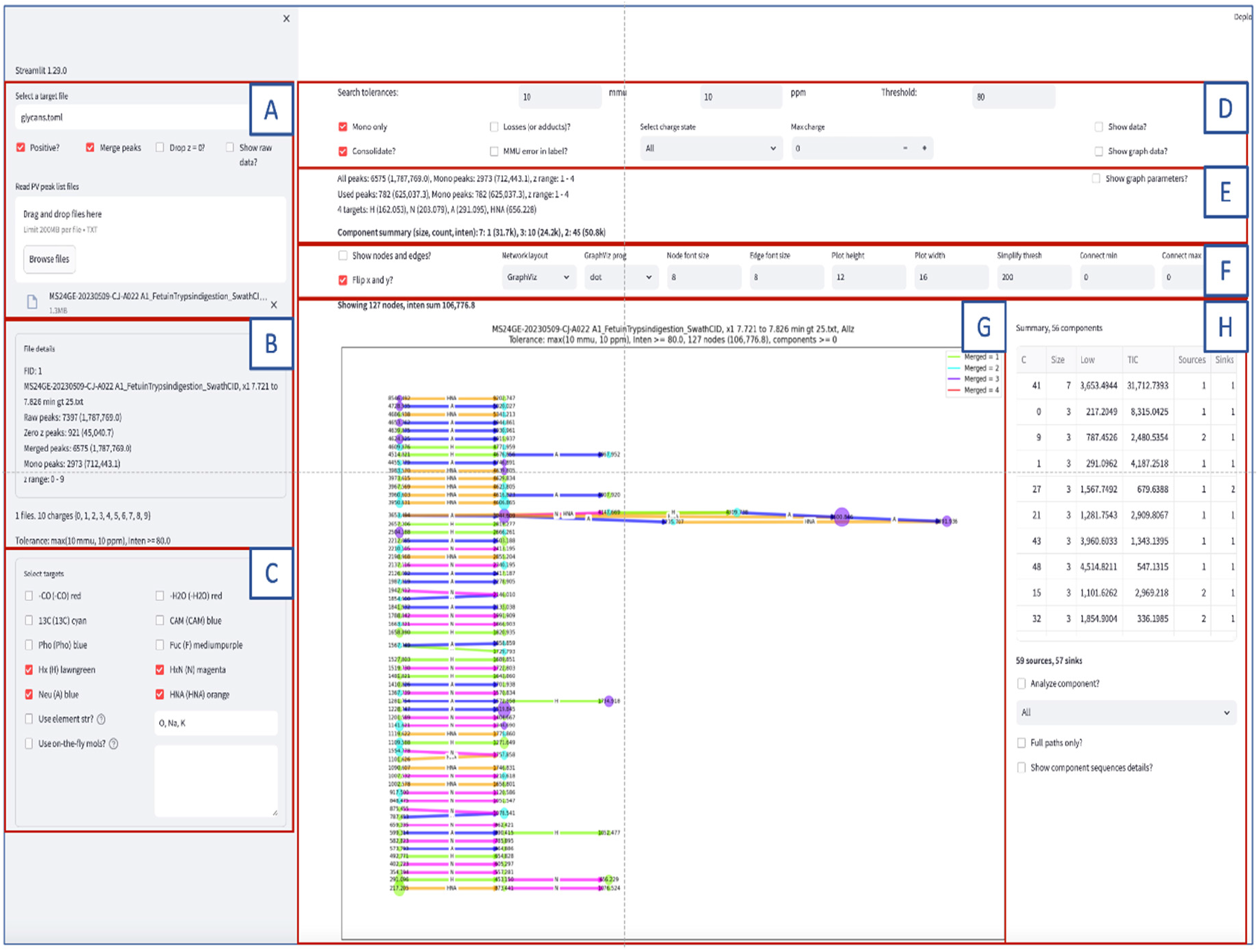
User interface. The sidebar contains A) Area for file selection and low-level parameter definition, B) File contents description, C) Target molecule definition area. The main area of the UI contains D) A region for commonly used processing parameters and controls, E) A summary of the peaks in the file (with intensity in brackets) and those used and/or monoisotopic, F) A region for graph control parameters, G) The main graph, H) A summary of the components present and controls for further exploring one or all components.

**Figure 3:**
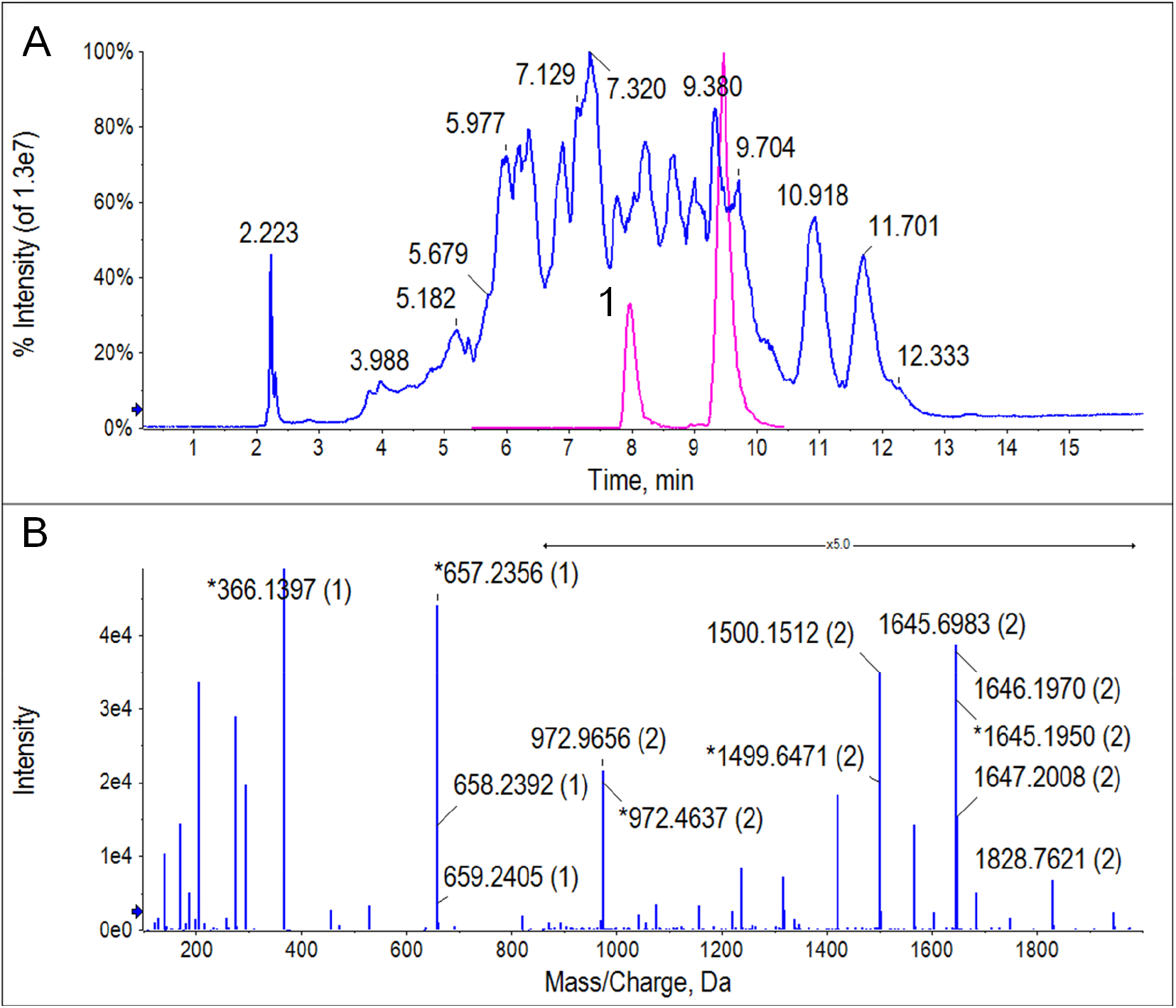
A) LC-MS/MS analysis of digested Fetuin, blue trace TIC, magenta trace XIC A, B) DDA-CID spectrum of peak 1 at RT = 7.9 min.

**Figure 4:**
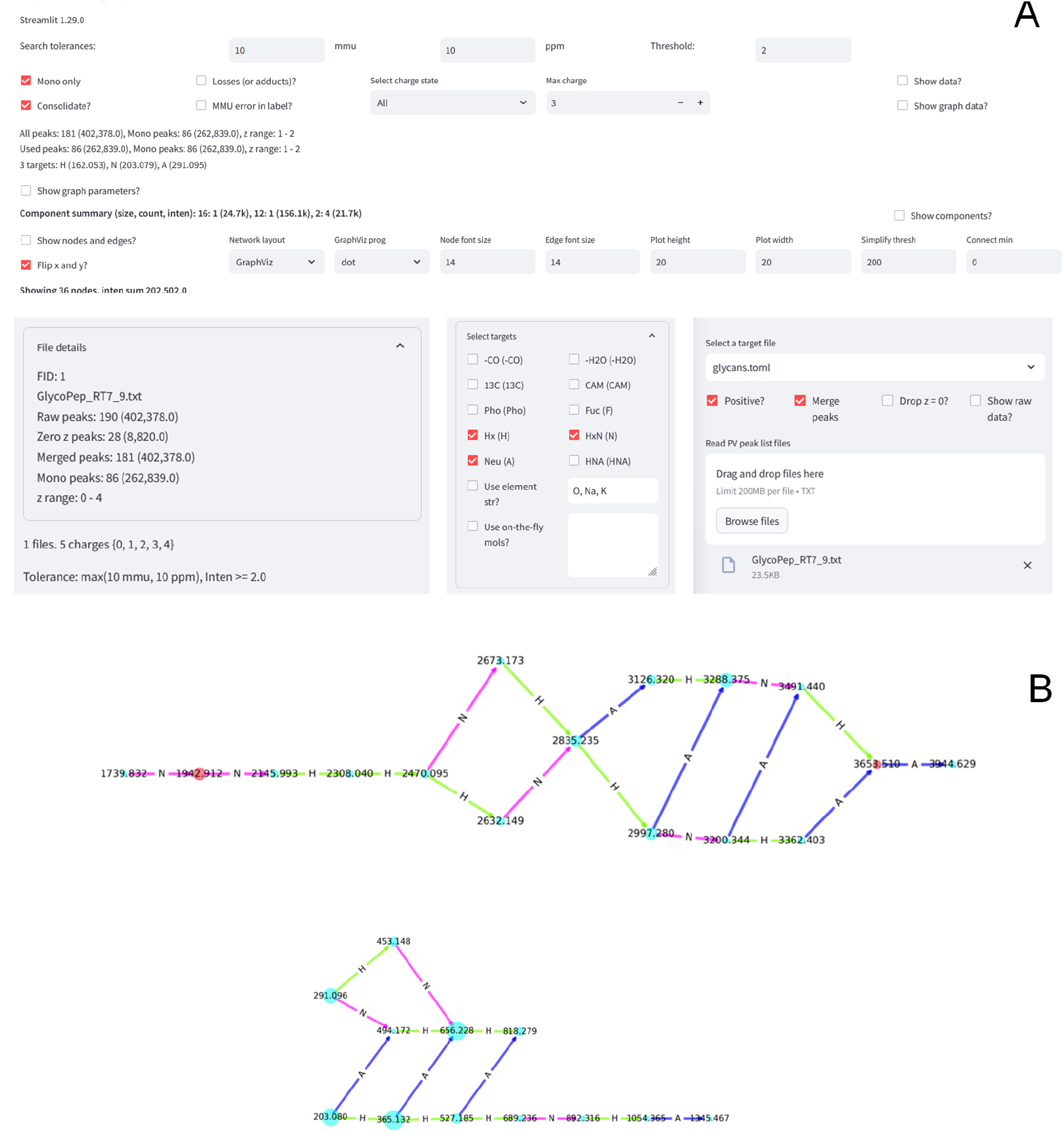
A) GlycanToGraphs settings for the analysis of MS/MS spectrum GlycoPep_RT7_9.txt and B) Output, the edges (arrows) indicate the linkages corresponding to the sugar masses H, N, A (glycans); the size of the symbols (here circles) reflects the intensity of the signals. The mass at 1739.832 corresponds to the mass of a peptide (P) and the mass at 3944.629 to P+H5N4A2

## 7. Conclusions

GlycansToGraphs is a python software tool developed for the unbiased visualization and filtering of complex relationships/sequences in the MS or MS/MS spectra of glycopeptides. The software requires only a spectrum text file and does not use protein or glycan databases. Mass differences are user defined and can correspond to any type of chemical modification, e.g. metal adducts (M, Na, Ca, K), solvent clusters. Homo-or hetero-multimers can be included by specifying their masses as arbitrary ‘on-the-fly’ molecules. The approach is not limited to the analyses of glycopeptides but can used in other applications, for example: other protein modifications, drug metabolism by hydroxyl or glucuronides, etc., or the study of complex adduct patterns.

## Author Contributions

Conceptualization: R.B., C.J., G.H.., Acquisition: J.J.., Software: R.B. Supervision: G.H., Visualization: J.L., J.U., L.T., D.B., Writing—Original Draft Preparation: R.B., C.J, G.H.., Writing—Review & Editing: R.B., CJ, G.H..

## Funding

This work was supported by the Swiss National Science Foundation, Grant 200021_192306.

## Conflict of Interest Statement

The authors declare no competing interests.

## Software download

The software can be downloaded at: https://doi.org/10.26037/yareta:beczz67s75d5xhk3xbjrzovipi

## Supplemental Figure

**Supplementary Figure S1.**
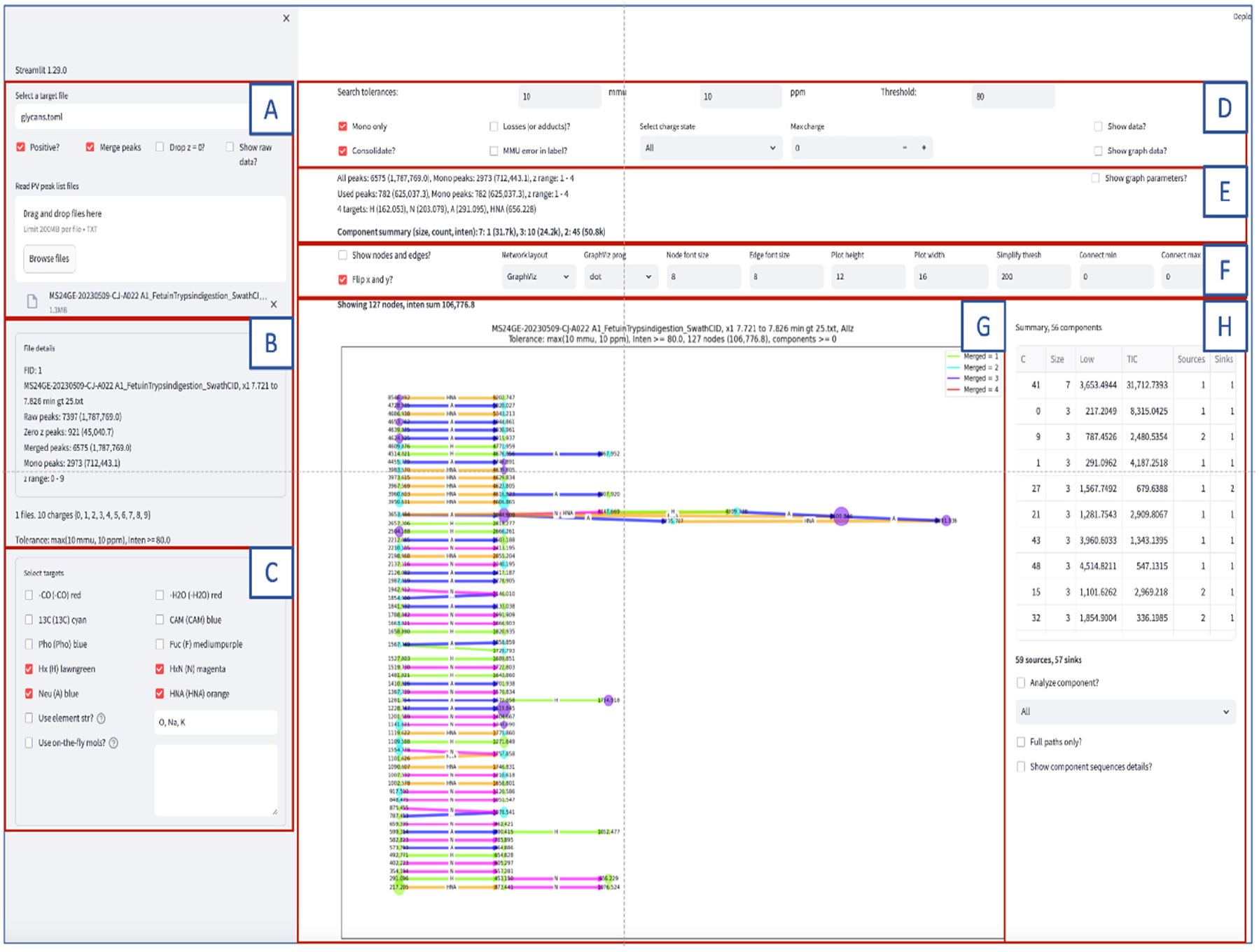
User interface details

The sidebar, which can be hidden to provide more space for the graph display, contains:

- Panel A: Collapsible area for file selection and low level parameter definition.

If ‘Merge peaks’ is true, masses are merged using the parameters defines in box D and ‘Z’ reflects the number of peaks merged (i.e. from different charge states).

Peak lists exported from PeakView can contain peaks with charge 0, indicating that there was no evidence of isotopes up to the maximum charge specified in the PV options. These peaks can be ignored or treated as singly charged.

‘Show raw data’ generates a table containing the raw peaks (m/z, intensity), charge and assigned mass. This can be useful for verifying the m/z values contributing to an observed mass.

- Panel B: File contents description.

A collapsible area that displays the peak details for the selected file(s)

- Panel C: Target molecule definition area

Molecules defined in the TOML configuration file are described by name, symbol and colour; the latter two are used to label edges in the main graph.

Elements can be added as a string and separated by commas. The defined elements are shown when the mouse is over the ‘?’ character and use the ‘effective adduct mass’, i.e. the mass delta is calculated considering the oxidation state so, for example, Fe can be entered as Fe2 (Fe II) or Fe3 (Fe III); most elements use the most common oxidation state.

Arbitrary masses can be entered separated by semicolons and may, optionally, include a symbol and/or a colour, i.e. the following are all valid: 146.056; 146.056,,DMSA; 146.056, red; 146.056, red, DMSA. This allows observed or expected masses to be quickly added to determine the effect on the network. Black is used if the colour is not specified and a symbol of the form M0, M1.. is generated if one is not specified.

The main display area contains:

- <u>Panel D</u>: A region for commonly used processing parameters and controls
- <u>Panel E:</u> A summary of the peaks in the file (with intensity in brackets) and those that are used and/or monoisotopic.

A summary of the size distribution of the network components as: size (number of nodes): count (number of components of this size) and summed intensity in thousands (k) in brackets

- Panel F: A region for graph control parameters

The layout is determined by the algorithm selected with ‘Network layout’ and, if GraphViz is selected, ‘GraphViz prog’. For the latter we have found ‘dot’ and ‘neato’ to be most useful.

‘Flip x and y’ causes the components to be drawn horizontally.

If the number of nodes displayed exceeds the ‘Simplify thresh’ the graph is simplified by decreasing the mass precision, hiding the edge labels and decreasing the width of the edges.

‘Connect min’ and ‘Connect max’ control the components displayed by specifying the size range (see box H). If ‘Connect max’ is 0, the maximum size is used.

- Panel 2: The main graph.

The legend describes the node colour and reflects the number of peaks (charge values) merged if ‘Merge peaks’ is true (box A) otherwise the charge (i.e. z in m/z).

Edges are drawn in the target colour indicated in box C and labelled with the corresponding symbol.

- Panel H: A summary of the components present and controls for further exploring one or all components. See section 5.6 ‘Component Analysis’ for more details

The processing workflow is summarized in Figure 1.

## Notes

### Competing Interest Statement

The authors have declared no competing interest.

